# Causal role of frontocentral beta oscillation in comprehending linguistic communicative functions

**DOI:** 10.1101/2024.04.03.587760

**Authors:** Wenshuo Chang, Xiaoxi Zhao, Lihui Wang, Xiaolin Zhou

## Abstract

Linguistic communication is often considered as an action serving the function of conveying the speaker’s goal to the addressee. Although neuroimaging studies have suggested a role of the motor system in comprehending communicative functions, the underlying mechanism is yet to be specified. Here, by 2 EEG experiments and a tACS experiment, we demonstrate that the frontocentral beta oscillation, which represents action states, plays a crucial part in linguistic communication understanding. Participants read scripts involving 2 interlocutors and rated the interlocutors’ attitudes. Each script included a critical sentence said by the speaker expressing a context-dependent function of either promise, request, or reply to the addressee’s query. These functions were behaviorally discriminated, with higher addressee’s will rating for the promise than for the reply and higher speaker’s will rating for the request than for the reply. EEG multivariate analyses showed that different communicative functions were represented by different patterns of frontocentral beta activity but not by patterns of alpha activity. Further tACS results showed that, relative to alpha tACS and sham stimulation, beta tACS improved the predictability of communicative functions of request or reply, as measured by the speaker’s will rating. These results convergently suggest a causal role of the frontocentral beta activities in comprehending linguistic communications.

**Highlights:**

- The frontocentral beta activity patterns discriminate communicative functions.
- The frontocentral beta activity plays a causal role in comprehending the functions.
- The comprehension could be subserved by the mental simulation of the communication.
- Our findings connect the idea of embedded semantics with the speech act theory.

## Introduction

Linguistic communication often involves 2 interlocutors, a speaker and an addressee. As suggested by prominent linguistic theories such as Sprachspiel (Wittgenstein, 1953) and speech act theory (Austin, 1975; Searle, 1969), the speaker’s utterance often conveys a certain communicative function rather than simply describe an objective situation. For example, when a speaker says “It is hot here” in an airtight room, the speaker’s utterance can function as a request to the addressee for turning on the air conditioning. Thus, linguistic communication can be viewed as an action serving the function of achieving the speaker’s goal (Austin, 1975; Searle, 1969; Wittgenstein, 1953).

Depending on the context, the speaker’s same utterance can serve a different communicative function, such as commissive, directive, assertive, declaration, or expressive (Searle, 1969). The present study focused on the first 3 categories. By commissives (e.g., promising, assuring), the speaker commits themselves to conduct a task that usually favors the addressee. In contrast, by directives (e.g., requesting, ordering), the speaker commits the addressee to conduct a task that usually favors the speaker. For example, by saying “I will revise your thesis” as a promise, a thesis advisor conveys a goal to help a student to improve the thesis. By saying “please revise my thesis” as a request, a student conveys a goal to seek help in improving the thesis from the advisor. While commissives and directives involve the interlocutors’ attitudes toward preferences concerning the tasks, assertives (e.g., replying to a query, stating a situation) are descriptions of situations that can be irrelevant to the interlocutors’ attitudes. For example, by saying “I revised the student’s thesis.” to reply a colleague’s query “What did you do this morning?”, a thesis advisor simply describes an event. Thus, the speaker’s utterance conveying distinct communicative functions in different situations also conveys their distinct attitude towards the current situation or task (Chang et al., 2022; Pérez Hernández, 2001).

As several theories consider linguistic communication as a kind of actions, neuroimaging studies indeed observed the involvement of the neural substrates for motor action not only in speech production/perception engaging motor programing (Wilson et al., 2004) but also in comprehending linguistic communicative functions conveyed by specific contents (Chang et al., 2022; Egorova et al., 2016; van Ackeren et al., 2012). For example, in our recent functional magnetic resonance imaging (fMRI) study (Chang et al., 2022), we found that the distinct communicative functions, such as promising, requesting, and replying, can be discriminated by the multivariate activity patterns in the premotor cortex. Moreover, these multivariate activities also contained the information about the interlocutors’ attitudes. Furthermore, patients with lesion in premotor cortex showed impaired comprehension of this information (Chang et al., 2022). These results demonstrated the role of the premotor cortex in processing communicative functions although the underlying mechanism remains to be specified. Given that the premotor cortex has been widely observed to be engaged during motor action processes with or without actual motor implementation (Hardwick et al., 2018), these findings suggest that the premotor cortex may subserve processes related to ones’ experience of actions (this kind of processes is usually termed as mental simulation, Savaki & Raos, 2019), and this finding leads to the hypothesis that the premotor cortex is involved in representing or simulating the interlocuters’ interaction based on ones’ communicative experience.

Similar to the findings on the premotor cortex, rhythmic brain activity approximately between 15 and 25 Hz (i.e., beta oscillation) in the motor system has been observed in motor action processes even without motor output (e.g., motor imagery, Brinkman et al., 2014; Ménoret et al., 2015; Ménoret et al., 2014). According to the “status quo” assumption, beta power varies with changing action states, with decreased power when an action is initiated and increased power when an action is inhibited (Engel & Fries, 2010). The role of beta oscillation in representing action state in the absence of motor implementation is supported by studies on motor imagery, showing that the beta power varied with the conditions of imagined actions, such as task demand (Brinkman et al., 2014), goals, and number of performers of actions (Ménoret et al., 2015).

Moreover, beta oscillations are also observed during the processing of visually or auditorily presented linguistic materials describing action semantics (for a review, see Weiss & Mueller, 2012). For example, just as observing manual actions would decrease frontocentral beta power, listening to sentences describing actions, either concrete (e.g., “Now I cut the bread”) or abstract (e.g., “I have drawn the consequence”), elicit similar decreases of frontocentral beta power (Moreno et al., 2013; Schaller et al., 2017). In an extension of action semantics, another EEG experiment showed that oscillatory activities within 11-18 Hz (beta band extending to high-alpha band) varied with the context-dependent communicative functions conveyed by the speaker’s sentences (Gisladottir et al., 2018). However, it is not clear yet whether the beta oscillation is causally involved in the processing of information concerning the potential action conveyed by linguistic communicative function.

To address this issue, we assessed beta oscillations during the processing of context-dependent communicative functions. We conducted 2 EEG experiments to test whether beta activity patterns contain information of communicative functions, and a transcranial alternating current stimulation (tACS) experiment to examine the causal role of beta oscillations in comprehending communicative functions.

## Experiments 1A and 1B: EEG study

### Methods

#### Participants

Forty-two native Chinese speakers (18 females, mean age = 22 years, standard deviation = 2.64, range: [18, 28]) participated in Experiment 1A, and 41 (13 females, mean age = 22 years, standard deviation = 2.58, range: [18, 29]) participated in Experiment 1B^1^. Two participants in Experiment 1A and 3 participants in Experiment 1B were excluded from data analyses due to noisy EEG recordings. Their remaining useful EEG epochs after preprocessing were lower than 60% in one or more experimental conditions. All the participants were recruited from universities in Beijing. They had normal or corrected-to-normal vision and had no known history of neurological or psychiatric disorders. Written informed consent was obtained from each participant prior to the experiment. The experiments were conducted in accordance with the Declaration of Helsinki and was approved by the Committee for Protecting Human and Animal Subjects of the School of Psychological and Cognitive Sciences at Peking University (No. #2019-10-04).

#### Design and materials

In Experiments 1A and 1B, participants were instructed to silently read scripts, each containing a critical sentence said by the speaker conveying a communicative function. All scripts were presented in written form, avoiding the potential impacts of prosodic information embedded in speech on the understanding of communicative function. While the critical sentence was presented as a whole in Experiment 1A, it was presented segment-by-segment in a rapid serial visual presentation (RSVP) mode in Experiment 1B to mitigate the potential influence of eye movements on EEG recording. Eye movements were recorded in Experiment 1A, and their potential confounding impacts on the EEG recording of processing the critical sentence were addressed during data analysis.

The same sets of scripts from our previous lesion study (Chang et al., 2022) were used in Experiments 1A and 1B. These scripts described daily-life communicative scenarios. Each script consisted of a context, a pre-critical sentence, and a critical sentence. Depending on the context, the critical sentence could have a different communicative function, which defined the 4 experimental conditions: *Promise*, *Reply-1*, *Request*, and *Reply-2* (see Table 1 for exemplars). The *Reply-1* and *Reply-2* served as control conditions, respectively, for the critical *Promise* and *Request* conditions. They simply described events without conveying specific interlocutor’s attitude.

**Table 1.** English translation and original Chinese version of the experimental scripts.

| Communicative function | Context | Precritical sentence | Critical sentence |
| --- | --- | --- | --- |
| *Promise* | Xiaoli was assigned to conduct data analysis for the survey this week. But Xiaoli was too busy to analyze the data. His colleague Wuyang had more spare time this week. And they were communicating with each other.<br>小李这周需要分析一批问卷调查数据, 但小李最近比较忙, 同事吴洋这周比较空闲, 于是两人进行了交谈。 | Wuyang said to Xiaoli,<br>吴洋对小李说: | “I will analyze the survey data this week.”<br>“这周我负责分析问卷数据。” |
| *Reply-1* | Wuyang was assigned to conduct data analysis for the survey this week. His colleague Xiaoli was recording the work memos, and wondering Wuyang's assignment of this week. And they were communicating with each other.<br>吴洋这周需要分析一批问卷调查数据, 同事小李负责记录工作安排, 想知道吴洋这周的任务, 于是两人进行了交谈。 | Wuyang said to Xiaoli,<br>吴洋对小李说: | “I will analyze the survey data this week.”<br>“这周我负责分析问卷数据。” |
| *Request* | Wuyang was assigned to conduct data analysis for the survey this week. But Wuyang was too busy to analyze the data. His colleague Xiaoli had more spare time this week. And they were communicating with each other.<br>吴洋这周需要分析一批问卷调查数据, 但吴洋最近比较忙, 同事小李这周比较空闲, 于是两人进行了交谈。 | Wuyang said to Xiaoli,<br>吴洋对小李说: | “I will analyze the survey data this week.”<br>“这周你负责分析问卷数据。” |
| *Reply-2* | Xiaoli was assigned to conduct data analysis for the survey. His colleague Wuyang recorded the work memos. Xiaoli wanted to confirm his assignment of this week. And they were communicating with each other.<br>小李这周需要分析一批问卷调查数据, 同事吴洋负责记录工作安排, 小李想确认自己这周的任务, 于是两人进行了交谈。 | Wuyang said to Xiaoli,<br>吴洋对小李说: | “I will analyze the survey data this week.”<br>“这周你负责分析问卷数据。” |

There were 84 quadruplets of scripts in total. As shown in Table 1, each quadruplet comprised 4 scripts corresponding respectively to the 4 conditions. Each script began with the context of an interpersonal interaction involving 2 interlocutors, a speaker and an addressee, followed by the sentence “and they were communicating with each other”. Subsequently, there were a pre-critical sentence “A (i.e., the speaker’s name) said to B (i.e., the addressee’s name)” and the final critical sentence. The critical sentence was said by the speaker to convey a communicative function and had a structure of “temporal subject + the first/second pronominal subject + predicate”. For the example in Table 1: “这周 (this week) + 我/你 (I/you) + 负责分析问卷数据 (will analyze the survey data)”. The *Reply-1* and *Reply-2* were used as controls for the *Promise* and *Request* respectively. Both these controls had the critical sentences being assertive sentences said by the speaker to reply the addressee’s queries. The critical sentences were varied only by the pronominal subjects, with the first-person pronoun for the *Promise* and *Reply-1* and the second-person pronoun for the *Request* and *Reply-2*.

The scripts were assigned into 4 experimental lists using a Latin-square procedure. Each list included 84 scripts (21 scripts for each condition) and was subdivided into 4 sections, with 5 or 6 scripts from each condition. Each section of scripts corresponded to a section of the main experimental task. In the task, each participant read a list of scripts in a pseudo-randomized order. They were required to complete 3 or 4 behavioral measures related to comprehension after reading each script.

#### Apparatus

The apparatus in Experiments 1A and 1B were the same except that an eye tracker was used in Experiment 1A.

##### Stimulus presentation

The tasks were programmed using *Psychtoolbox* (Brainard, 1997) in MATLAB, and the stimuli, in black (RGB: 0, 0, 0) against a gray background (RGB: 180, 180, 180), were presented on a Display++ LCD monitor (screen area: 70×40 cm^2^, resolution: 1920×1080 pixels, Cambridge Research Systems, UK). The distance between the computer screen and participants’ eyes was set at 60 cm using a chinrest. Each character on the screen occupied approximately 1 degree of visual angle.

##### Eye tracking

In Experiment 1A, participants’ eye movements were tracked by a desktop-mounted EyeLink 2000 system (SR Research, Canada) at a sampling rate of 2000 Hz. The eye movements recording and calibrations were carried out monocularly based on the right eye while the actual viewing was binocular.

##### EEG system

For both Experiments 1A and 1B, participants’ EEG activities were recorded with 64 Ag/AgCl electrodes mounted in an antiCap (Brain Products, Germany, see Figure 1A for the channel montage based on the 10-20 system). The electrooculogram (EOG) was recorded by 2 electrodes: vertical EOG was recorded by an electrode placed above the right eye, and horizontal EOG was recorded by an electrode placed at the outer canthus of the right eye. All electrode impedances were kept below 10 kΩ. The EEG and EOG recordings were amplified by the BrainAmps (Brain Products, Germany) with the AFz channel as a ground and FCz channel as an online reference, and digitized online at a sampling rate of 500 Hz. (EEG data of one participant were online sampled at 1000 Hz and offline down-sampled to 500 Hz to keep consistency across participants.) An online notch filter was used on EEG data to remove power line noise at 50 Hz.

**Figure 1.**
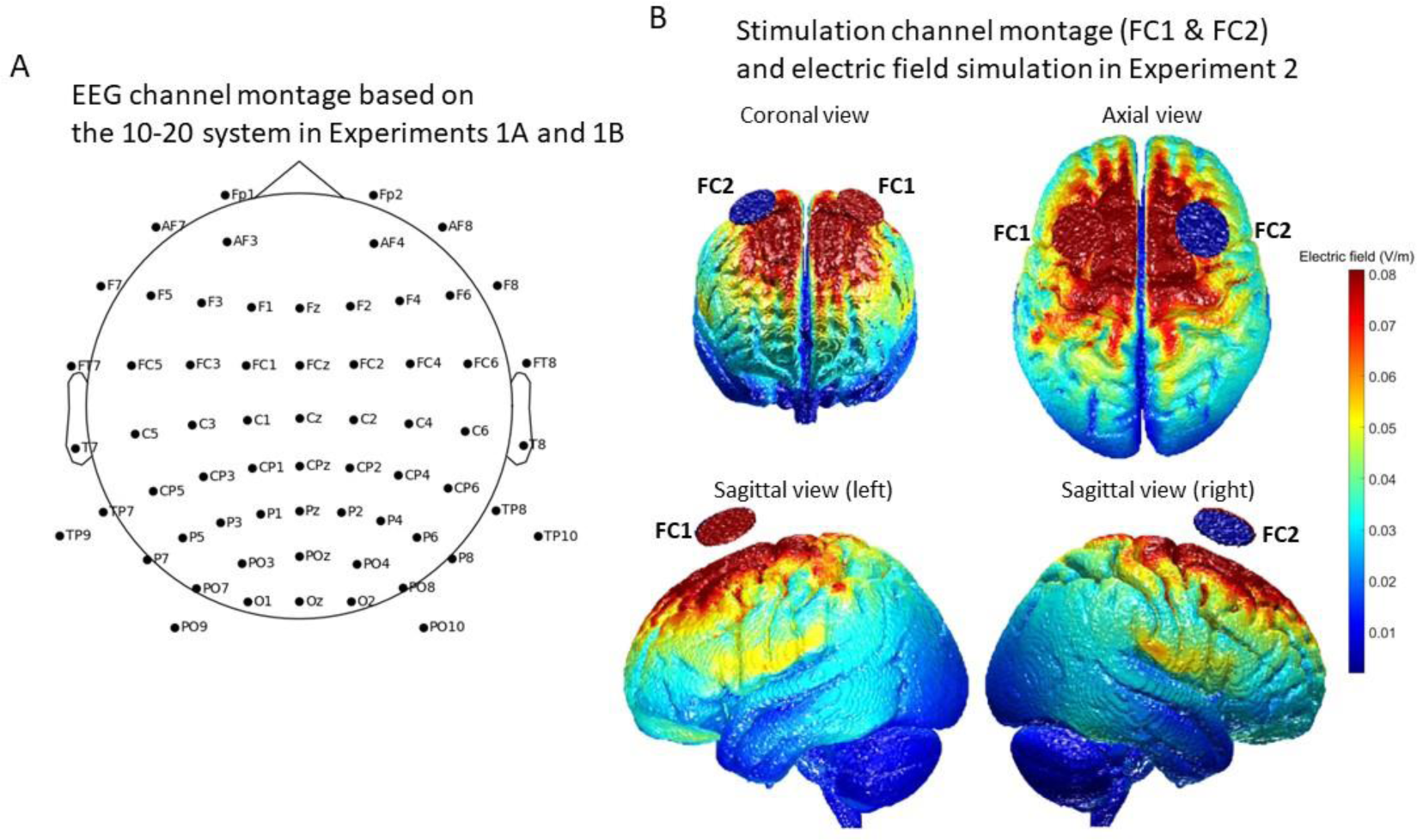
Channel montages. (**A**) EEG channel montage based on the 10-20 system in Experiments 1A and 1B. (**B**) Stimulation channel montage (FC1 and FC2) and electric field simulation of the stimulation in Experiment 2. The red and blue discs illustrate the FC1 and FC2 channels respectively.

#### Procedure

##### Experiment 1A

Data of EEG and eye movements were simultaneously recorded. The 4 sections of the main task started after 10 practice trials. At the beginning of each session, participants completed a calibration procedure with a nine-point grid for eye tracking, followed by a warm-up trial. As illustrated in Figure 2A, each trial started with a fixation cross at the center for a jittered duration of 1 to 3 s, followed by a cross at the upper left site where the first character of the context would locate. This fixation was presented for 0.5 s to direct participants’ eye gaze. Then the context sentence was presented as a whole for 10 s and followed by a cross presented for 0.5 s at the upper left site, where the first character of the pre-critical sentence would locate. Then the pre-critical sentence was presented as a whole for 1 s. After the offset of the pre-critical sentence, a cross was presented below the first character of the pre-critical sentence, where the first character of the critical sentence would locate. This cross was presented for 0.5 s together with the pre-critical sentence, followed by the critical sentence for 2s. The critical sentence was inside double quotes and presented together with the pre-critical sentence. This presentation of the critical sentence was designed to allow natural reading.

**Figure 2.**
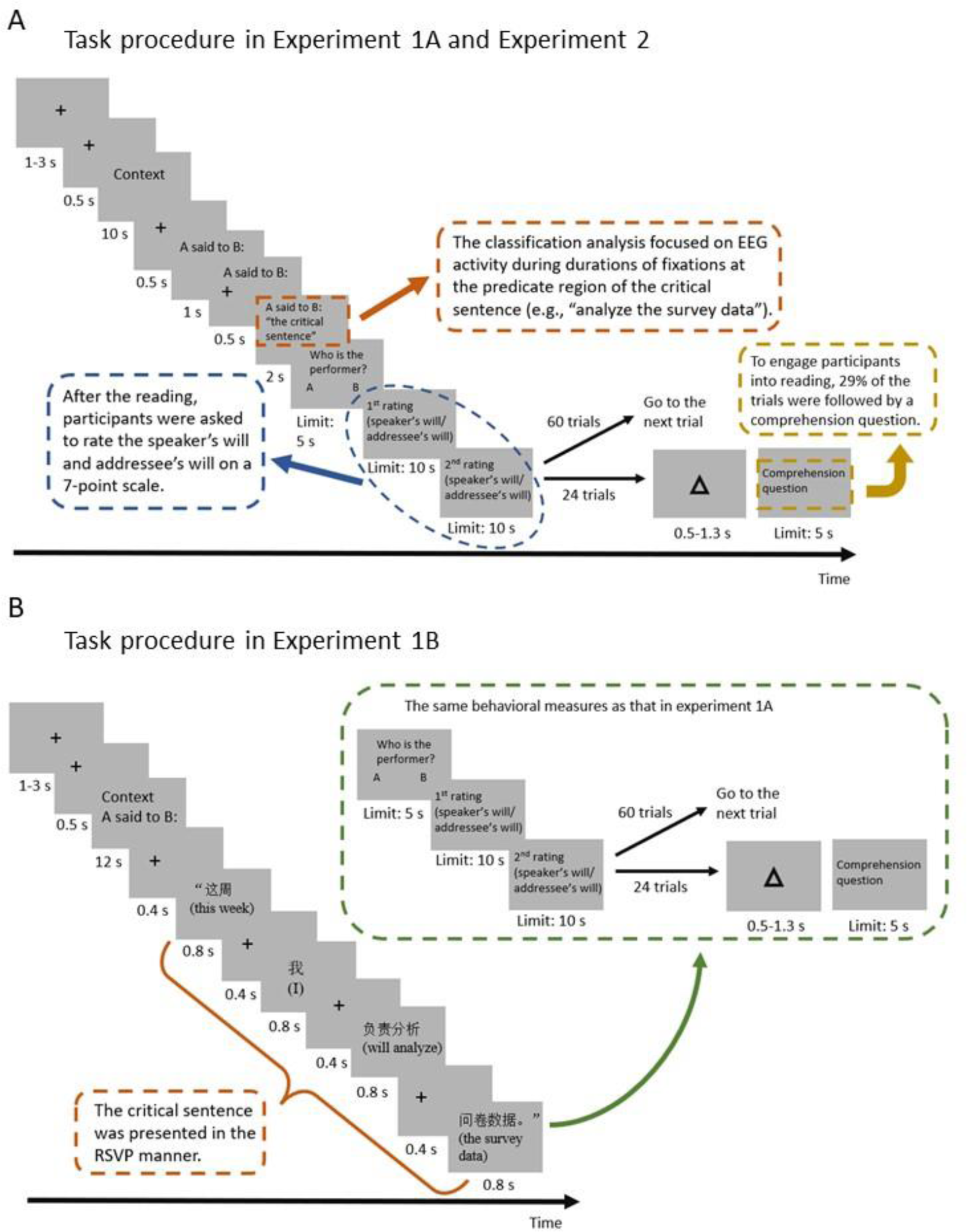
Task procedures. (**A**) The procedure in Experiments 1A and 2. (**B**) The procedure in Experiment 1B. The English translations in parentheses are presented for illustration and were not presented in the experiment.

After each script, participants were asked to perform 3 or 4 behavioral measures, following our previous study (Chang et al., 2022). First, they were asked to judge which of the speaker and the addressee would be an expected performer of the task described in the critical sentence (i.e., the performer judgment). The two names were presented at the lower left and lower right of the screen respectively, and the mapping between the two locations and the two names was randomized across trials. Participants had to press the button on the corresponding side to choose either of the names within 5 s. Second, they were asked to rate the speaker’s will and the addressee’s will, on a 7-point scale within 10 s. The order of the ratings on the speaker’s will and the addressee’s will was random. To engage participants into the reading, a Yes/No comprehension questions regarding the scripts followed the ratings in 29% of trials (i.e., catch trials, 6 catch trials were included for each condition). In each catch trial, a triangle was presented at the screen center after the ratings and before the comprehension question, and this triangle was presented for a jitter duration of 0.5 to 1.5s. Chinese words “yes” and “no” were presented at the left bottom and the right bottom of the screen respectively for half of participants, and the reversed mapping of these words and locations was used for the other half. Participants had to answer the question by pressing the corresponding button within 5 s.

##### Experiment 1B

During EEG recording, participants were instructed to perform the same task as in Experiment 1A, with the only difference being that the critical sentences were presented segment by segment in a RSVP mode to mitigate the potential confounding effect of eye movements on EEG recording. As shown in Figure 2B, in the RSVP, the critical sentence was presented sequentially in 4 segments: the temporal subject with a left quotation mark (e.g., “这周” means “this week”), the pronominal subject (e.g, “我” means “I”), the first part of the predicate, henceforth the predicate-1 (e.g., “ 负责分析” means “will analyze”), and the second part of the predicate, henceforth the predicate-2 with a right quotation mark (e.g., “问卷数据” means “the survey data”). Each segment was preceded by a cross fixation presented for 0.4 s and was presented for 0.8 s.

As communicative functions are associated with the speaker/addressee’s will regarding the task described by the predicate, our EEG analyses focused on the activities during the reading of the predicate. Specifically, in Experiment 1A, we focused on the EEG activity during the duration of fixations at the predicate area (e.g., “负责分析问卷数据”/“will analyze the survey data”); in Experiment 1B, we focused on EEG activities during presentation of 3 interested segments: the predicate-1 (“负责分析”/“will analyze”), cross fixation following the predicate-1, and predicate-2 (“问卷数据”/“the survey data”).

#### Speaker/addressee’s will rating analysis

To examine the extent to which the ratings of the speaker’s will and addressee’s will predicted communicative functions, we fitted Bayesian hierarchical logistic models using *Stan* (Carpenter et al., 2017) in R for 2 paired predictions, “*Promise* vs. *Reply-1*” and “*Request* vs. *Reply-2*”, respectively. Each model was fitted with the logit function and included the communicative function as the response variable and the speaker’s will rating and addressee’s will rating as predictors, as shown in Equation 1.

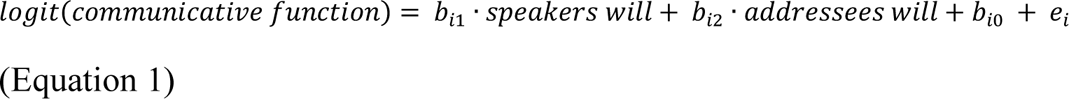

In each model, the parameters, including the slopes of the predictors and the intercept, were estimated at the participant-level to assess individual variance for each participant, and were also estimated at the group-level to assess population effects. Each group-level parameter had a normal prior distribution, which had a mean with a prior of *Normal*(0, 100) and a standard deviation with a prior of *Cauchy*(0, 5). Each participant-level parameter had a normal prior distribution, which had a mean equaling to the mean of the group-level distribution and a standard deviation with a prior of *Cauchy*(0, 5). Equation 1 shows these parameters: for the *i*th participant, *b*_*i*1_, *b*_*i*2_, *b*_*i*0_, and *e*_*i*_ indicate the slopes of the speaker’s will and addressee’s will, the intercept, and the residual respectively. The posterior distribution of each parameter was sampled using 4 Monte-Carlo Markov Chains (MCMCs), each comprising 10000 iterations. The initial half of each chain was discarded as warm-up samples. Mean posterior estimate and 95% credible interval (*CrI*) were reported to evaluate the effect of each parameter. All posterior estimates reported had *Ȓ* -values lower than 1.01, indicating their convergence (Gelman & Rubin, 1992). An estimated effect with 95% *CrI* excluding 0 would be interpreted as statistically significant.

#### EEG preprocessing

EEG data were preprocessed using the *MNE* package (Gramfort et al., 2014) in python, and the same preprocessing method was applied to both experiments. To recover the FCz signal, which was used as the online reference, the EEG data were re-referenced with average amplitude of all 63 channels (including the FCz). To ensure the quality of independent component analysis (ICA), the data were band-pass filtered between 1 and 100 Hz to remove noise for fitting ICA model with extended Infomax algorithm (Lee et al., 1999; Winkler et al., 2015). This band-pass filter was applied only to the process involved the ICA. For further analysis, the data were high-pass filtered above 0.1 Hz to remove low frequency line noise, avoiding the potential degradation of the EEG data due to filter method (Tanner et al., 2015).

The filtered data were segmented into epochs containing EEG responses to the predicate of the critical sentence. In Experiment 1A, EEG and eye-tracking data were coregistered using the *EYE-EEG* plugin (Dimigen et al., 2011) in the *EEGLAB* toolbox (Delorme & Makeig, 2004). Then epochs were segmented from -1.5 to 2.5 s relative to the onset of the first fixation at the predicate area (trials with no fixation at the predicate were excluded). In Experiment 1B, epochs were segmented from -16.3 to 4.6 s relative to the onset of the predicate-1. These relatively long epochs were to ensure the quality for fitting ICA model.

ICA model was fitted based on the band-pass filtered epochs to decompose vertical and horizontal EOG artifacts, electrocardiogram (ECG) artifact, and channel noise. Artifact components were identified by visual inspection and by putative component types classified by the *ICALabel* package (Li et al., 2022; Pion-Tonachini et al., 2019). The artifacts were subtracted from the 0.1 Hz high-pass filtered epochs (Winkler et al., 2015). Outlier epochs were identified as those involving peak-to-peak amplitudes exceeding 100 μV after additional low-pass filtering below 30 Hz, and they were excluded from data analysis. Finally, in the epochs with 0.1 Hz high-pass filter but without a low-pass filter, outliers were removed to obtain artifact-free epochs for further analyses.

#### Time-frequency transform

Time-frequency transforms of the EEG epochs were conducted by band-pass filters and Hilbert transforms for alpha (8-12 Hz) and beta (18-25 Hz) bands. For each frequency band, band-pass filtering, Hilbert transform, and baseline correction were applied to the epochs. In Experiment 1A, the baseline interval of -100-0 ms relative to the onset of the first fixation at the predicate was chosen. In Experiment 1B, the baseline interval of -100-0 ms relative to the onset of the interested segment was chosen. Following baseline correction, the signal was squared to obtain oscillatory power.

#### Region of interest (ROI)

Two regions of interest (ROIs) were defined: (1) frontocentral region, comprising channels of F7, F5, F3, F1, Fz, F2, F4, F6, F8, FC5, FC1, FC2, FC6, FC3, FC4, FCz, C5, C3, C1, Cz, C2, C4, and C6; (2) parietooccipital region, comprising channels of P7, P3, Pz, P4, P8, P5, P1, P2, P6, PO9, PO7, PO3, POz, PO4, PO8, PO10, O1, Oz, and O2. Centroparietal (CP) channels lying between the two regions were excluded from analyses.

#### Multivariate pattern classification (MVPC)

MVPCs were conducted on the oscillatory power to show if the oscillations contain distinct information of the communicative functions. Two paired classifications were performed, “*Promise* vs. *Reply-1*” and “*Request* vs. *Reply-2*”. These classifications were implemented using the *scikit-learn* package (Pedregosa et al., 2011) in python for each frequency band, each ROI, and each participant.

In Experiment 1A, for each epoch (trial), the oscillatory power during durations of all fixations within the predicate area was extracted and were averaged across time for the 63 channels, yielding a vector of 63 data points. For each channel and each communicative function (condition), the vectors of data were averaged across trials within the same section. In Experiment 1B, the oscillatory power was extracted and averaged across time for each of the 3 interested segments. Subsequently, the power for each segment was averaged across trials within the same section. These calculations yielded a 4 (section) × 63 (channel) × 2 (“*Promise* and *Reply-1*” or “*Request* and *Reply-2*”) matrix for each participant and each paired classification in Experiment 1A and a 4 (section) × 63 (channel) × 2 (“*Promise* and *Reply-1*” or “*Request* and *Reply-2*”) matrix for each participant, each paired classification, and each segment in Experiment 1B.

A support vector machine (SVM) classifier was used to train and cross-validate each paired classification, “*Promise* vs. *Reply-1*” or “*Request* vs. *Reply-2*”. The data was partitioned into 4 folds, each corresponding to a section. The “leave-one-fold-out” cross-validation was applied with 3 folds included in the training set and the remaining fold included in the test set and was repeated 4 times to obtain an average classification accuracy. The discrimination performance of each paired classification was evaluated by an across-participant average area under receiver-operator curve (AUC).

Statistical significance of AUCs was assessed by permutation tests. To form a null distribution, for each participant, individual-level null AUCs were obtained by permutating the communicative function labels during the training and cross-validation. This permutation was repeated 1000 times for each paired classification to get the individual-level null AUC set for each participant. Then the group-level null AUCs were obtained by randomly sampling AUCs from the individual-level null sets (with replacement) and by averaging these individual-level null AUCs. This procedure was repeated 100000 times, resulting in a null distribution of group-level AUCs. The probability of the observed group-level AUC in the null distribution of group-level AUCs was calculated. Upper-tailed *p*-value was used to assess statistical significance with false discovery rate (FDR) correction for multiple comparisons.

The MVPCs were performed on the beta activity for our research purpose, and also performed on the alpha activity as a control to show the specificity of the beta frequency.

## Results and Discussion

Participants had mean accuracies of 94% (*SD* = 8%) and 92% (*SD* = 10%) for performer judgement in Experiment 1A and Experiment 1B, respectively. They had average accuracies of 85% (*SD* = 9%) and 84% (*SD* = 10%) for comprehension questions in Experiment 1A and Experiment 1B, respectively.

Bayesian hierarchical logistic models were fitted to predict communicative functions by the ratings of the speaker’s will and the addressee’s will. Results of model estimates showed the same pattern in Experiments 1A and 1B, as shown in Figure 3A and 3B. For the “*Promise* vs. *Reply-1*” models, communicative functions were significantly predicted by the addressee’s will rating (*b* = 2.45, 95% *CrI*: [2.05, 2.92] in Experiment 1A; *b* = 2.71, 95% *CrI*: [2.07, 3.48] in Experiment 1B), with a higher addressee’s will rating for *Promise* than for *Reply-1* but no effect in the speaker’s will rating. In contrast, for the “*Request* vs. *Reply-2*” models, communicative functions were significantly predicted by the speaker’s will rating (*b* = 1.74, 95% *CrI*: [1.45, 2.07] in Experiment 1A; *b* = 1.94, 95% *CrI*: [1.6, 2.33] in Experiment 1B), with a higher speaker’s will rating for *Request* than for *Reply-2*. In this model, communicative functions were also significantly predicted by the addressee’s will (*b* = -1.59, 95% *CrI*: [-2.04, -1.25] in Experiment 1A; *b* = -1.22, 95% *CrI*: [-1.61, -0.9] in Experiment 1B), with a lower rating for *Request* than for *Reply-2*. These results suggested that the same critical sentence could contain different social information that distinguishes communicative functions. Specifically, the communicative function of *Promise* is predicted by the will of the addressee, who benefits from the fulfillment of the promised task, whereas the communicative function of *Request* is predicted by the will of the speaker, who benefits from the fulfillment of the requested task.

**Figure 3.**
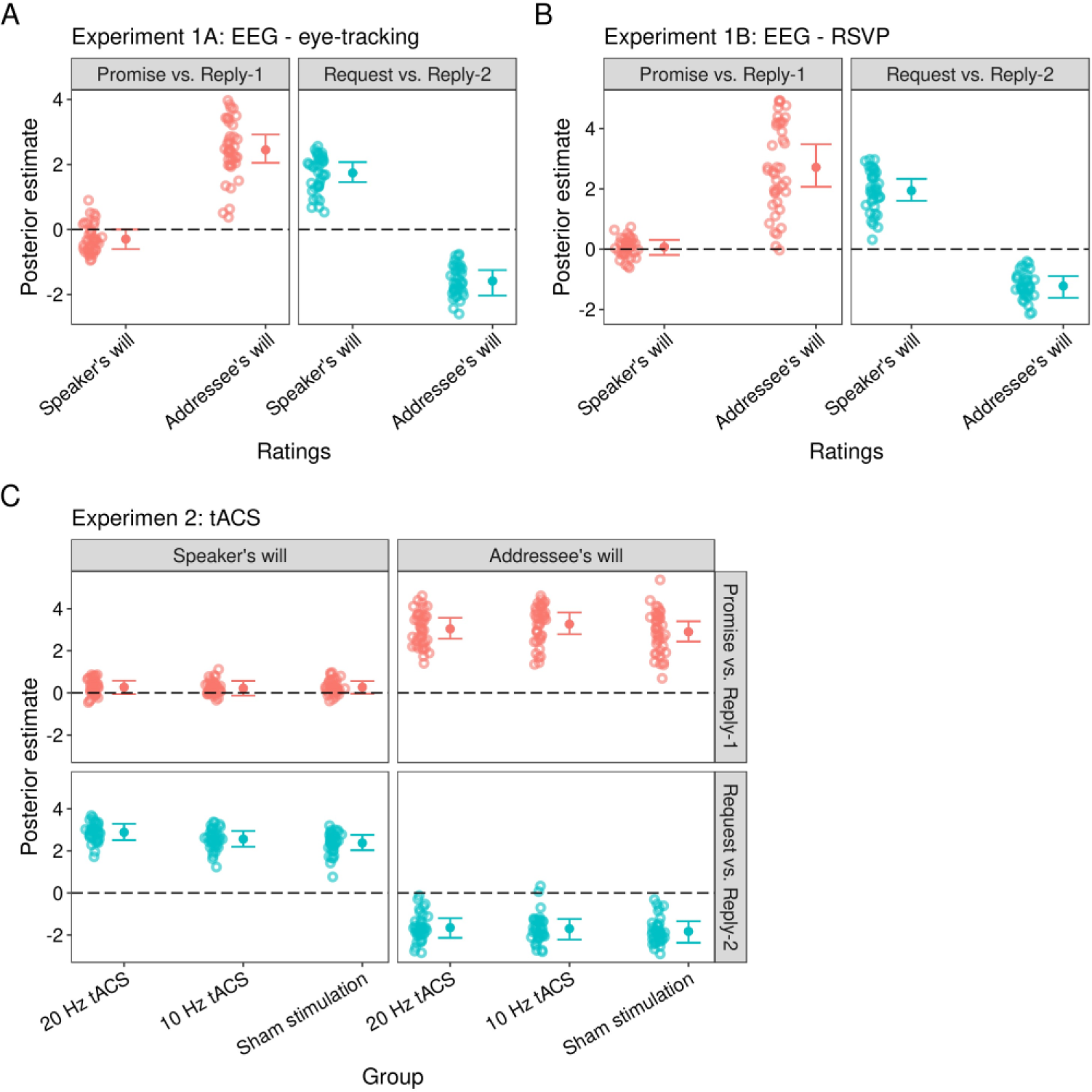
Results of Bayesian hierarchical logistic models. (**A** and **B**) Results in Experiments 1A and 1B, respectively. The horizontal axes indicate the predictors, the speaker’s will rating and the addressee’s will rating. (**C**) Results in Experiment 2. The left panel shows results of the speaker’s will rating and the right panel shows results of the addressee’s will rating. The horizontal axes indicate the stimulation conditions, 20 Hz tACS, 10 Hz tACS, and sham stimulation. (**A**-**C**) The vertical axes indicate posterior slope estimates of these predictors. The solid circles indicate mean group-level posterior estimates. The error bars show 95% *CrI*s of group-level estimates. The crowded circular rings indicate participant-level estimates. The black dashed lines show the position where posterior estimate equals to 0.

To examine if beta and alpha activity patterns contained the information of communicative functions, we conducted MVPCs to decode communicative functions from the activity patterns in the frontocentral and parietooccipital regions, respectively. The classification performances were indexed by areas under receiver-operator curve (AUCs).

### Experiment 1A

For beta activity in the frontocentral region, as shown in Figure 4A, AUCs of both “*Promise* vs. *Reply-1*” and “*Request* vs. *Reply-2*” classifications were significantly above chance-level (AUC = 53%, *p* = 0.035; AUC = 54%, *p* = 0.002, respectively). No significant result was showed for alpha activity (*p*-values > 0.08).

**Figure 4.**
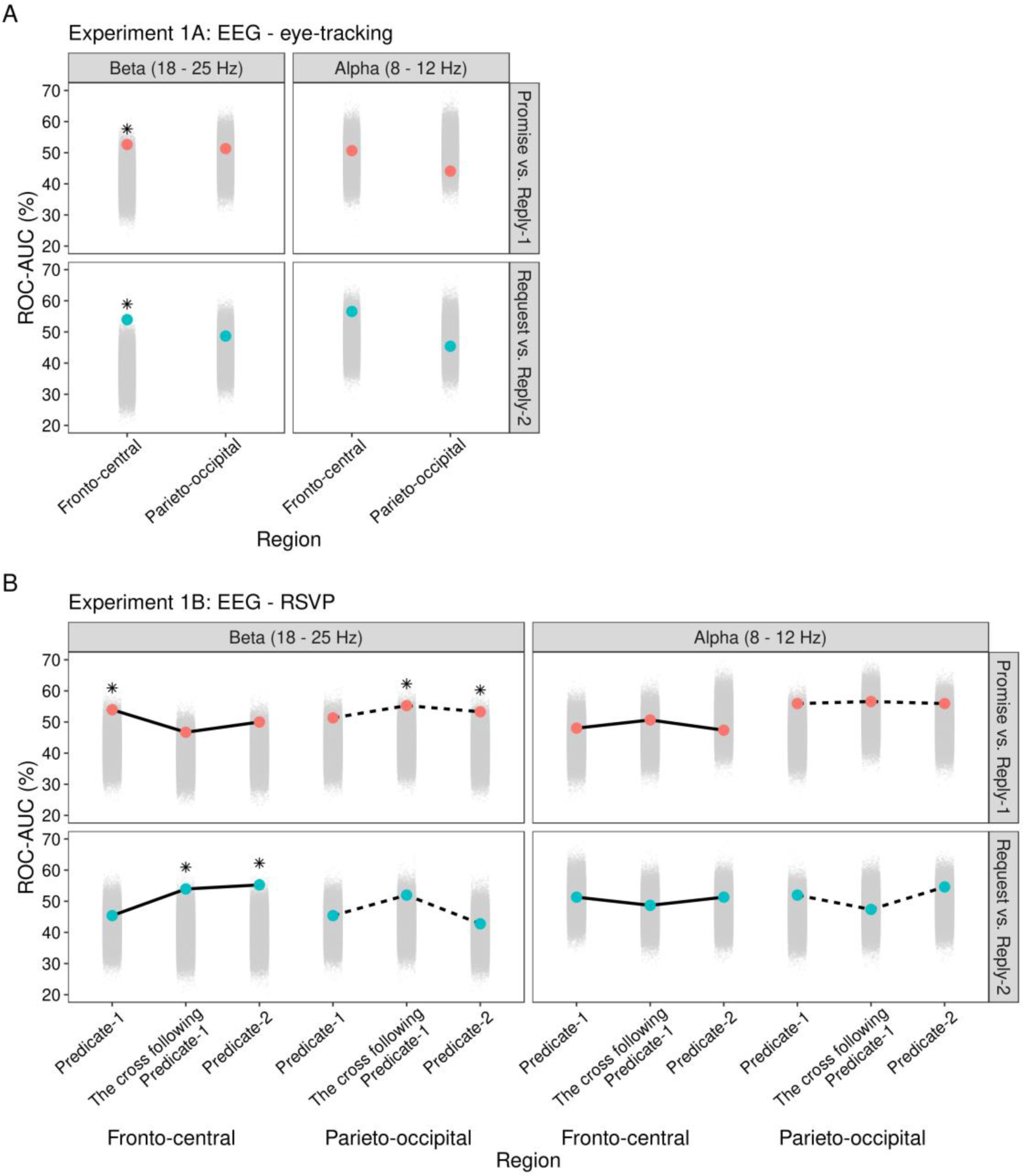
MVPC results. (**A**) Results in Experiment 1A. The horizontal axes indicate the regions. (**B**) Results in Experiment 1B. The horizontal axes indicate the segments of the critical sentence for the regions respectively. (**A** and **B**) The upper panel shows results for the “*Promise* vs. *Reply-1*” classification and the lower panel shows results for the “*Request* vs. *Repy-2*” classification. The left panel shows results for beta oscillation and the right panel shows results for alpha oscillation. The vertical axes indicate ROC-AUC values. The solid circles indicate the mean ROC-AUC across 50 cross-validations. The crowded gray solid circles indicate the permuted distributions for AUCs. The black stars indicate significant results for the permutation tests.

### Experiment 1B

For beta activity in the frontocentral region, as shown in Figure 4B, the AUC of the “*Promise* vs. *Reply-1*” classification was significantly above chance-level during the presentation of the predicate-1 (AUC = 54%, *p* = 0.035), and AUCs of the “*Request* vs. *Reply-2*” classification were significantly above chance-level following the predicate-1 (AUC = 54%, *p* = 0.024) and during the presentation of the predicate-2 (AUC = 55%, *p* = 0.018). For beta activity in the parietooccipital region, AUCs of the “*Promise* vs. *Reply-1*” classification were also significantly above chance-level following the predicate-1 (AUC = 55%, *p* = 0.035) and during the presentation of the predicate-2 (AUC = 53%, *p* = 0.035). No significant result was showed for the alpha activity (*p*-values > 0.05).

Thus, the results of MVPCs in both experiments suggested that the beta activity patterns in the frontocentral region robustly represent the information of communicative functions, whereas alpha activity shows no such distinctive patterns.

## Experiment 2: tACS study

### Methods

#### Participants

One hundred and twenty native Chinese speakers, all from universities in Shanghai, participated the tACS experiment. Written informed consent and safety screening survey for transcranial electrical stimulation (tES) were obtained from each participant prior to the experiment. All participants reported no risk factor for tES, had normal or corrected-to-normal vision, and had no known history of neurological or psychiatric disorders. Each participant was assigned to one of 3 stimulation groups, resulting in 42 participants in the 20 Hz tACS group, 40 participants in the 10 Hz tACS group, and 40 participants in the sham stimulation group. In the 10 Hz tACS group, 2 participants’ data were excluded from the analyses. One of them was excluded due to his approximately chance-level accuracy of performer judgement (49%), and the other was because the stimulation condition was mistakenly set as sham in the last two sections. Hence, each group had an equal sample size (40 participants per group, see Table 2 for their demographic information). The 3 groups had comparable ages as suggested by the Bayes factor favoring H_0_ against H_1_ for one-way analysis of variance (ANOVA) (*BF*_01_ = 11.4). The experiment was conducted in accordance with the Declaration of Helsinki and was approved by the Committee for Protecting Human and Animal Subjects of the School of Psychological and Cognitive Sciences at Peking University (No. #2021-07-02).

**Table 2.** Demographic information of participants and descriptive statistics of the questionnaire on the experience during tES in Experiment 2.

| Group | Sex | Age<br>(years) | Performer<br>judgement<br>accuracy<br>(%) | Comprehension<br>question<br>accuracy (%) | Sensation rating on 5-point scale (0 - 4) |  |  |  |  |  |  | Guess of the stimulation<br>condition (number count) |  |  |
| --- | --- | --- | --- | --- | --- | --- | --- | --- | --- | --- | --- | --- | --- | --- |
|  |  |  |  |  | Itching | Pain | Burning | Warmth | Pinching | Metallic | Fatigue | Real | Sham | I don't<br>know |
| 20 Hz | 23 females/ | 23 (3) | 91 (9) | 88 (8) | 0.7 | 0.15 | 0.13 | 0.05 | 0.08 | 0 | 0.95 | 22 | 4 | 14 |
| tACS | 17 males | [18,30] |  |  | (0.85) | (0.36) | (0.33) | (0.22) | (0.27) | (0) | (1.01) |  |  |  |
| 10 Hz | 22 females/ | 23 (2) | 93 (8) | 86 (8) | 0.5 | 0.3 | 0.25 | 0.35 | 0.38 | 0 | 1.15 | 18 | 8 | 14 |
| tACS | 18 males | [20,30] |  |  | (0.64) | (0.72) | (0.54) | (0.58) | (0.59) | (0) | (1.1) |  |  |  |
| Sham | 21 females/ | 23 (3) | 94 (7) | 86 (8) | 0.68 | 0.33 | 0.25 | 0.25 | 0.15 | 0.05 | 1.2 | 13 | 6 | 21 |
| stimulation | 19 males | [18,29] |  |  | (0.8) | (0.62) | (0.54) | (0.49) | (0.48) | (0.32) | (1.02) |  |  |  |

Demographic information and descriptive statistics are presented for the 3 groups respectively. The means, standard deviations (in round brackets) of ages, the two types of accuracies, and the 7 sensation ratings, the range of ages (in square brackets), and number counts of guesses at stimulation conditions are presented.

#### Apparatus

Transcranial alternating current stimulation (tACS) was administered using the Starstim 8 system with the Instrument Controller (NIC) software (Neuroelectrics, Spain). Two circular sponge-based electrodes (8 cm^2^) were connected with the current control box by wires, and were respectively placed at the FC1 and FC2 channels on the EEG-cap. The electric field distribution of this channel montage in the brain was simulated using the computational model of the Realistic vOlumetric Approach by the *ROAST* toolbox (Huang et al., 2019), showing that the medial premotor cortex can be covered in the area with the highest electric field (Figure 1B). The sponge-based electrodes were soaked in saline solution (NaCl) to keep the impedance below 10 kΩ. The impedance was monitored based on the CMS and DRL channels placed over the right mastoid during the stimulation. The stimulation would be automatically interrupted by the system once the impedance exceeds 20 kΩ for safety.

#### Procedure

Prior to the experiment, participants were exposed to 10 Hz tACS for 110 s (including 30 s ramp-up and 20 s ramp-down) at 1 mA and subsequently at 2 mA (peak-to-peak). This was to familiarize the participants with tACS. The 10 Hz tACS was chosen because it was reported to induce stronger sensation than the beta frequency tACS (Fertonani et al., 2015). None of the participants reported intolerance of the stimulation.

The same practice task and main experimental task as in Experiment 1A were presented to participants using a laptop (Dell G3 3590). Following the familiarization, participants completed the practice task without stimulation and proceeded to engage in the 4 sections of the main task with simultaneous tACS. Each participant was assigned to one of 3 stimulation groups: 20 Hz (beta band) tACS, 10 Hz (alpha band) tACS, and sham stimulation. During each section of the task, the stimulation was administered at 2 mA (peak-to-peak) intensity and had a sinusoidal waveform of 0^°^relative phase without direct current offset. For the 20 Hz tACS and 10 Hz tACS, the stimulation during each section was started with a 30 s ramp-up, lasted 13 min, and ended with a 20 s ramp-down. For the sham stimulation, an alternating current at 10 Hz (to keep consistent with the frequency used for familiarization), was started with a 30 s ramp-up, followed by a period of no stimulation for 13 min, and ended with a 20 s ramp-down. For each section, the ramp-up and ramp-down phases occurred when no experimental stimuli were presented.

After the experiment, participants completed a Chinese version of the questionnaire on the experience during tES (Fertonani et al., 2015). In the questionnaire, they were asked to rate their perceived intensities of 7 listed sensations on 5-point scales (from 0, indicating no sensation, to 4, indicating strong sensation) and to guess the stimulation condition as either “real”, “sham”, or “I don’t know”. The descriptive statistics of these data are shown in Table 2. The low mean ratings of sensations indicated that participants did not have uncomfortable sensations during stimulation. To compare questionnaire responses between groups, Bayes factors favoring H_0_ against H_1_ (*BF*_01_) for contingency tables with independent multinomial sampling were computed (Gunel & Dickey, 1974). The Bayes factors showed that the 7 sensation ratings (*BF*_01_ range: [3.08, 284.92]) and the number count of each alternative guess of stimulation condition (*BF*_01_ = 11.33) were comparable between the groups.

#### Accuracies of the performer judgments and comprehension questions

Frequentist and Bayesian one-way ANOVAs were performed to compare accuracies of the performer judgments and comprehension questions between the 3 groups. We report *F*-statistics, degrees of freedom (*df*s), effect size estimates (*η*_*p*_^2^), *p*-values, and Bayes factors ( *BF*_01_) in supporting H_0_ against H_1_ in the Results and Discussion section.

#### Speaker/addressee’s will rating analysis

To examine the extent to which the participants’ ratings of the speaker’s will and addressee’s will could predict communicative functions and to compare the model estimates between the 3 groups, we fitted Bayesian hierarchical logistic models with logit function for “*Promise* vs. *Reply-1*” and “*Request* vs. *Reply-2*” respectively. To reduce the potential confounding effect of participants’ reading comprehension abilities, we adopted the data screening criterion in our previous lesion study (Chang et al., 2022) to include only the trials with correct performer judgements. This procedure left us with 92% of the data. Each hierarchical logistic model included communicative functions as the response variable, the ratings of the speaker’s will and the addressee’s will as predictors, and each participant’s guess of the stimulation condition as a covariate whose variability should be regressed out (see Equation 2). The participant’s guess of the stimulation condition was coded as 1, if “real” was chose, -1, if “sham” was chose, or 0, if “I don’t know” was chose.

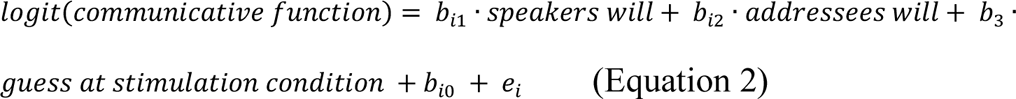

In each model, the slopes of the speaker’s will and the addressee’s will and the intercept were estimated at the group-level and participant-level, respectively. The group-level estimation assessed average effects of predictors for each group. The participant-level estimation assessed effects of predictors for each participant and provided individual observations for between-groups comparisons. Equation 2 shows the model parameters: for the *i*th participant, *b*_*i*1_, *b*_*i*2_, *b*_*i*0_, and *e*_*i*_ have the same meanings as Equation 1. The slope of the guess of the stimulation condition was shown as *b*_3_ and had a normal distribution with a mean having a prior of *Normal*(0, 100) and a standard deviation having a prior of *Cauchy*(0, 5). The settings of priors for other parameters and the model-fitting method were the same as that of the model of Equation 1. All posterior estimates reported have *Ȓ*-values lower than 1.01. An estimated effect with 95% *CrI* excluding 0 would be interpreted as statistically significant.

To examine the tACS effect on the predictability of communicative functions from the ratings, we compared the participant-level slope estimates of the 2 ratings between the 3 groups using Frequentist and Bayesian one-way ANOVAs and conducted post-hoc comparisons using Frequentist and Bayesian *t*-tests in R. These slope estimates were considered as indices of participants’ capability of comprehending communicative functions. We report *F*-statistics, *t*-statistics, *df*s, effect size estimates (Cohen’s *d* for *t*-tests, and *η*_*p*_^2^ for ANOVAs), *p*-values, and Bayes factors (*BF*_10_) in supporting H_1_ against H_0_ in the Results. We interpreted the strength of evidence based on Bayes factors following Andraszewicz and colleagues’ suggestion (see Table 1 in Andraszewicz et al., 2015).

## Results and Discussion

### Accuracies of the performer judgments and comprehension questions

All 3 groups had mean accuracies of the performer judgements above 91% and of the comprehension questions above 85%. Results of one-way ANOVAs showed that the 3 groups had comparable accuracies of the performer judgements (*F*_(2,117)_ = 0.94, *η*_*p*_^2^ = 0.02, *p* = 0.395, *BF*_01_ = 5.83) and the comprehension questions (*F*_(2,117)_ = 0.84, *η*_*p*_^2^ = 0.01, *p* = 0.433, *BF*_01_ = 6.3). These results indicated that the between-group tACS effects regarding communicative functions cannot be attributed to general differences in sentence comprehension.

### Results of the speaker/addressee’s will rating analyses

We fitted Bayesian hierarchical logistic models, including the ratings of the speaker’s will and the addressee’s will as predictors, to predict communicative functions. Results are shown in Figure 3C.

For the “*Promise* vs. *Reply-1*” model, the group-level slope estimates of the 3 groups showed that communicative functions were significantly predicted by the addressee’s will ratings (20 Hz tACS: *b* = 3.04, 95% *CrI*: [2.57, 3.57]; 10 Hz tACS: *b* = 3.26, 95% *CrI*: [2.79, 3.81]; sham stimulation: *b* = 2.9, 95% *CrI*: [2.44, 3.4]), indicating a higher rating for *Promise* than for *Reply-1*. One-way ANOVA on the participant-level slope estimates of the addressee’s will rating showed no significant between-group differences (*F*_(2,117)_ = 1.69, *η*_*p*_^2^ = 0.03, *p* = 0.189, *BF*_10_ = 0.32).

For the “*Request* vs. *Reply-2*” model, the group-level estimates of the 3 groups showed that communicative functions were significantly predicted by the speaker’s will ratings (20 Hz tACS: *b* = 2.88, 95% *CrI*: [2.51, 3.29]; 10 Hz tACS: *b* = 2.56, 95% *CrI*: [2.2, 2.95]; sham stimulation: *b* = 2.38, 95% *CrI*: [2.03, 2.76]) and the addressee’s will ratings (20 Hz tACS: *b* = -1.65, 95% *CrI*: [-2.13, -1.2]; 10 Hz tACS: *b* = -1.7, 95% *CrI*: [-2.21, -1.23]; sham stimulation: *b* = -1.83, 95% *CrI*: [-2.37, -1.34]), indicating higher speaker’s will rating and lower addressee’s will rating for *Request* than for *Reply-2*. One-way ANOVA on the participant-level slope estimates of the speaker’s will rating showed extreme evidence for between-group difference (*F*_(2,117)_ = 12.59, *η*_*p*_^2^ = 0.18, *p* < 0.001, *BF*_10_ = 1747.63). Post-hoc comparisons obtained extreme evidence for the higher slope estimates of the speaker’s will rating for the 20 Hz group than for either the 10 Hz group (2.88 vs. 2.56, *t*_(75.99)_ = 5.03, Cohen’s *d* = 1.13, *p* < 0.001, *BF*_10_ = 5043.09) or the sham stimulation group (2.88 vs. 2.37, *t*_(76.59)_ = 3.28, Cohen’s *d* = 0.73, *p* = 0.002, *BF*_10_ = 20.93), but no significant difference between the 10 Hz group and the sham stimulation group (2.56 vs. 2.37, *t*_(77.94)_ = 1.7, Cohen’s *d* = 0.38, *p* = 0.094, *BF*_10_ = 0.8). One-way ANOVA on the slope estimates of the addressee’s will rating showed no significant between-group difference (*F*_(2,117)_ = 0.86, *η*_*p*_^2^ = 0.01, *p*= 0.427, *BF*_10_ = 0.16).

## General Discussion

Across 3 experiments, we demonstrated causal roles of the frontocentral beta oscillation in processing linguistic communicative functions. In a replication of our previous study (Chang et al., 2022), we first showed that communicative functions were behaviorally distinguished by higher addressee‘s will rating for the *Promise* than for the *Reply-1*, and higher speaker’s will rating for the *Request* than for the *Reply-2*. In the MVPC results of the 2 EEG experiments, for both the “*Promise* vs. *Reply-1*” and “*Request* vs. *Reply-2*” classifications, classifiers based on the frontocentral beta activity patterns showed above-chance classifications of communicative functions regardless of the presentation manner of the scripts, whereas the results of the alpha oscillation showed no significant classification. The tACS experiment further showed that, in the “*Request* vs. *Reply-2*” model, the slope estimates of the speaker’s will rating were enhanced by 20 Hz (beta) tACS relative to either 10 Hz (alpha) tACS or sham stimulation over the frontocentral region covering the medial premotor cortex, suggesting a causal role of the frontocentral beta oscillation in comprehending linguistic communications. Collectively, these results support the hypothesis that the comprehension of linguistic communications involves the representations of the interlocutors’ interaction, echoing with our previous findings regarding the premotor cortex (Chang et al., 2022) and the theoretical view of considering linguistic communications as a kind of action (Austin, 1975; Searle, 1969; Wittgenstein, 1953).

Our findings advance the understanding of the cognitive functions in language processing subserved by beta oscillation in at least two aspects. First, in language processing, the frontocentral beta oscillations represent not only action semantics, as shown by previous studies (Moreno et al., 2013; Schaller et al., 2017; Weiss & Mueller, 2012), but also context-dependent communicative functions. Indeed, similar effects on beta oscillation has been reported by a previous EEG study (Gisladottir et al., 2018) which showed that, compared with sentences with either assertive or commissive functions, oscillatory activities within 11-18 Hz deceased for sentences with declination function. However, the authors did not differentiate activities of beta and alpha bands and did not differentiate activities in different EEG channels, rendering it difficult to interpret the specific role of frequency band and spatial region. By separately analyzing oscillatory activities within different frequency bands and regions, we showed that the beta activity pattern varies as a function of communicative function independently of semantic content of the critical sentence. This communicative function can be seen as a state of action, and the beta activity pattern reflects this function as part of its role in responding to action state, namely, the “status quo” assumption (Engel & Fries, 2010).

Second, the association of the frontocentral beta oscillation with comprehension of linguistic communication suggests that the experience-based processing of linguistic communications may involve the mental simulation of the communicative situations. Both EEG and tACS results on beta oscillation are consistent with previous observations on the variation of beta activity pattern in responding to action conditions (e.g., high or low task demand, or presence or absence of goal) during motor imagery (Brinkman et al., 2014; Ménoret et al., 2015). These general effects of the beta oscillatory fluctuation with various conditions could suggest that the comprehension of linguistic communications and motor imagery involve a common mechanism, and this mechanism could be the mental simulation of the encountered situation of action (Avenanti et al., 2007; Brinkman et al., 2014; Stadler et al., 2011).

Specifically, in comprehending linguistic communications, the comprehender likely mentally simulates the interlocutors’ interaction involving their attitudes toward the communicated content. This was evidenced by the enhanced beta tACS effect on the predictability of communicative functions based on the speaker’s will rating, and by the effect of premotor lesion on the speaker’s will and addressee’s will ratings in our previous study (Chang et al., 2022). To interpret the attitudes, the comprehender can take the perspective of the speaker, the initiator of the communication, and engage in a simulated situation and mentally build a model of the interactive situation between the interlocutors. The comprehender’s preference of taking the perspective of an action initiator over that of an action observer has been demonstrated during mentally simulating actions presented through linguistic materials (Brunyé et al., 2009) or videos of movements (Brattan et al., 2015). Information of the interlocutors’ attitudes then assists the comprehender to distinguish specific communicative functions. Such a process of understanding communicative functions is consistent with the idea of the idealized cognitive model (Lakoff, 1987; Pérez Hernández, 2001), which assumes that each communicative function has a prototypical model determining the interlocutors’ attitudes. Moreover, as suggested by the action prediction theory of communicative function (Tomasello, 2023), the simulated model can facilitate the comprehender to predict potential responsive action of the addressee. Take the request in Table 1 for an example, the comprehender can understand the speaker’s will to have the addressee’s help for data analysis and predict that the addressee would engage in the requested task. Such a prediction regarding communicative function is supported by a previous study showing that, during face-to-face communication, the activation of the speaker’s primary motor cortex is initiated before the onset of the speaker’s actual speech of requesting (Boux et al., 2021).

Findings in the current study contribute to our knowledge concerning sensorimotor representations in language processing. According to the mirror neuron hypothesis (Rizzolatti & Sinigaglia, 2016) and the embodied semantics (Bechtold et al., 2023; Della Putta, 2018), the mirror property of the motor system supports understanding action semantics by activating neural representations of relevant motor action (Hauk et al., 2004). The present results extend the embodied semantics to a more general sense by demonstrating that the frontocentral beta activity responds to the context-dependent social information beyond the semantic meaning of the speaker’s words. The motor system is involved not only in understanding semantic meanings of specific actions but in integrating contextual information. This argument is supported by studies on pragmatics (Feng et al., 2017; Feng et al., 2021) and by clinical observations (Li et al., 2023). For example, the functional connectivity between the medial premotor cortex and right temporo-parietal junction (TPJ), a brain region subserving the inference of others’ mental states, increased with context-dependent indirectness of replies (e.g., a sentence “Nowadays, people are really beginning to enjoy opera.” would be an indirect reply to “Do you think that the audience liked my opera performance?”), demonstrating a general role of the motor system in pragmatic inferencing (Feng et al., 2017).

To conclude, by using EEG and tACS techniques, we provided both correlational and causal evidence for the role of the frontocentral beta oscillation in comprehending linguistic communications. These results suggest that the beta oscillation in the motor system subserves the experience-based processes of linguistic communications. Our findings support the theoretical view that linguistic communication is represented as a kind of actions in the brain (Austin, 1975; Searle, 1969; Wittgenstein, 1953).

## Authors’ contributions

Chang, Wang, and Zhou designed the experiments. Chang and Zhao conducted the experiments. Chang analyzed the data. Chang, Wang, Zhao, and Zhou wrote the paper.

## Acknowledgements

We thank Ms. Mengqiao Deng, Mr. Ruiming Zhu, and Ms. Ping Ou at Shanghai Jiao Tong University for their assistance during data collection in the tACS experiment. The data analyses were conducted on the High-performance Computing Platform of Peking University.

## Data and code Availability

The raw data and code for data analyses can be accessed via https://osf.io/q4jfk/?view_only=afc114b7723c4a098a6af819d30e2099.

## Footnotes

1 As 11 participants' demographic information in Experiment 1B was lost due to storage system problem, the demographic statistics were based on the remaining 30 participants.

